# AarMIXTAs: key factors of T-shaped non-glandular trichome development in *Artemisia argyi*

**DOI:** 10.1101/2025.03.06.641095

**Authors:** Xinlian Chen, Duan Wu, Chunyu Li, Baosheng Liao, Qi Shen

## Abstract

Trichomes are necessary for stress resistance and metabolite synthesis in plant. T-shaped non-glandular trichomes (TSTs) as the main component of moxa floss, which are the specific characteristics of *A. argyi*. Here, we observed that TSTs were at an extremely high density on the lower surfaces in *A. argyi* leaves. The key transcription factors (TFs) involved in regulation of TSTs development of *A. argyi* were identified through RNA sequencing of TSTs and non-TSTs. The core regulatory genes, *AarMIXTA*s showed allelic amplification and functional differentiation compared with *A. annua*, might contribute development of TSTs. The overexpression of *AarMIXTA1.2* significantly increased the density of TSTs in transgenic Arabidopsis and may regulate trichome development by regulating the downstream genes, which played an important positively regulatory role in the development of TSTs. This study provides a good reference for the molecular mechanisms related to the development of TSTs and a theoretical basis for the high-yield moxa floss cultivation of *A. argyi*.

## Introduction

Trichomes are surface appendages differentiated from epidermal cells in the aerial part of plants (Ishida et al., 2008) . They exist on the surface of most Angiosperm, some Gymnosperm and Bryophyta plants, such as the fruit thorns of cucumbers, the fibers of cotton, and various tissues of plants (Han et al., 2022) . Trichomes are essential for plants to adapt to the environment and overcome biotic and abiotic stresses (Feng et al., 2021) . According to whether they can secrete secondary metabolites, they are usually divided into glandular secretory trichomes (GSTs) and non-glandular trichomes (NGTs) (Liu et al., 2017). NGTs do not have secretory or metabolic functions, but provide a physical barrier for plants, which can help plants reduce water transpiration, collect fog, enhance tolerance to abnormal temperatures, protect plants from ultraviolet radiation and herbivores, etc (Karabourniotis et al., 1992).

*Artemisia argyi* Lévl. et Vant is the perennial plant of Asteraceae family (Sciences et al., 1991) . Its dried leaves are the major medicinal materials in China. They have the effects of warming the meridians, stopping bleeding, dispelling cold and relieving pain; and externally removing dampness and relieving itching (Commission, 2020) . It is widely used in medicine, food, and culture. The germplasm resources of mugwort are distributed in China, Mongolia, North Korea, and cultivated in Japan (Sciences et al., 1991) . In China, the most famous *A. argyi* are in Qichun, Hubei (“Qi mugwort”) and Nanyang, Henan (“Nanyang mugwort”). In 2023, the comprehensive output value of mugwort industry have exceed ¥30 billion in China. Mugwort is widely used in treatment and health care as moxibustion therapy. Moxibustion, the traditional Chinese therapy with a history of thousands of years, uses burning moxa floss to warm the body surface or stimulate acupuncture points to prevent and treat diseases (Chang et al., 2012) . The dried leaves of *A. argyi* are used to make moxa floss, the primary material for moxibustion, which mainly consists of fibrous substances, particularly trichomes from *A. argyi* leaves. The therapeutic effect of moxibustion is predicted to be influenced by both the burning heat of moxa floss and the release of volatile substances from trichomes.

*A. argyi* have abundant NGTs, which are also referred to as T-shaped non-glandular trichomes (TSTs) based on their shape, especially on its lower epidermis (Cui et al., 2022) . Scanning electron microscopy revealed that TSTs of *A. argyi* exhibited unique characteristics, with particularly dense distribution on the lower epidermis and elongated to entangle with each other, which is the main tissue of moxa floss. The difference in the distribution of TSTs on the upper and lower surfaces of *A. argyi* leaves makes us propose that it is an ideal system to study the development of multicellular TSTs. Previous studies have confirmed that the yield and quality of moxa floss, directly affect the clinical effect of moxibustion, are closely related to the growth and trichome development.

The development of NGTs is usually studied using Arabidopsis rosette leaves or cotton fibers (Wang et al., 2019) . The NGTs of the two species are single-cell NGTs. As huge single epidermal cells, they are easy to analyze at the genetic, genomic and cell biological levels and have become a model system for studying cell development (Wang et al., 2019). However, there is no better research system for multicellular NGTs. The development of plant trichomes is regulated by a complex molecular network that has been well elucidated in Arabidopsis and *A. annua* (Yan et al., 2018) . The MYB-bHLH-WD1 complex acts as positive regulators, including GL1, GL3/EGL3, and TTG1, to activate downstream activators like GL2 and TTG2 (Yang & Ye, 2013) . Additionally, R2R3-MYB and HD-ZIP IV transcription factors (TFs) play essential roles in glandular trichome initiation in *A. annua* and tomato (Chalvin et al., 2020) . In 1994, Noda et al first cloned the *MIXTA* gene from *Antirrhinum majus* that controls flower color concentration and petal gloss (Noda et al., 1994) . *MIXTA*, belonging to subgroup 9 A (SBG9-A) of R2R3-MYB, is a gene that controls the shape of petal epidermal cells (Brockington et al., 2013) . Since then, *MIXTA*/*MIXTA-like* genes in more species have been found to be related to the regulation of plant epidermal cell differentiation, which in turn closely affects the development of trichomes (Wu et al., 2018; Zhao et al., 2020; Lashbrooke et al., 2015) .

Considering the biological traits of *A. argyi*, the developmental mechanism underlying trichome formation and regulation remain unclear. Here, we observed epidermis microscopic features of *A. argyi* leaves. The transcriptome of TSTs and non-TSTs of *A. argyi* leaves were sequenced to identify the key regulation factors and *AarMIXTA*s as core regulators attained further investigated for evolutionary relationships, expression differentiation, and functional verification. The research greatly enriched the development and regulation mechanism of plant trichomes, and provided a reliable theoretical basis for metabolic regulation to improve the yield and quality of *A. argyi*.

## Materials and Methods

### Preparation and observation of *A. argyi* for microscope

We selected two intact *A. argyi* leaves at the same leaf position growing well in the net house, and observed the distribution of trichomes on the upper and lower epidermis under the stereomicroscope. We selected 2-3 *A. argyi* leaves growing well in the net house. The leaves were cut out a square of 0.5 cm × 0.5 cm and placed in the center of a slide. Then we added water, covered with a cover glass and observed under a fluorescence microscope. We selected *A. argyi* that grew well in the net house, picked 2-3 leaves avoiding the main veins and cut into 0.4 cm×0.4 cm squares, immersed the cut leaves into 5% glutaraldehyde and placed them in the 4L refrigerator for two weeks. Then we took out the samples from the refrigerator and suck out the glutaraldehyde. The samples were rinsed for three times with 0.1 M PBS buffer (10 min/time), 30% ethanol, 50% ethanol, 70% ethanol, and 90% ethanol (10 min/time) in turn at room temperature. Finally, the samples were rinsed for three times with anhydrous ethanol (10 min/time). After washing, we used the critical point dryers to dry, mounting on stage, ion sputtering instrument to spray, and observed the GSTs and TSTs under the scanning electron microscope at different magnifications.

### RNA-seq and analysis

TSTs on the lower epidermis and non-TSTs of *A. argyi* leaves (the remaining of the leaves after removing TSTs) were manual separated for RNA-seq. Three biological replicates for each tissue. A RNAprep Pure Plant kit (TIANGEN, China) was used to extract total RNA, and 20 μg RNA was taken for reverse transcription to synthesize cDNA. RNA sequencing was performed on Illumina HiSeq X Ten platform. Reads from RNA-seq were aligned to the genome using HISAT2 and reconstructed transcripts by StringTie. The counts from featureCounts were used to calculate FPKM (fragments per kilobase of exon model per million mapped fragments). We performed cluster analysis, statistics on the number of expressed genes, statistics on the number and distribution of differentially expressed genes, and GO functional enrichment analysis on the six samples between TSTs and non-TSTs.

### WGCNA of transcriptome

To clarify more information of trichome development network, we download transcriptomic data of multiple tissues in *A. argyi* from NCBI (National Center for Biotechnology Information) (Table S1) to speculate expression trend with TPM (transcripts per million reads) and WGCNA (Weighted Correlation Network Analysis). We used R package WGCNA of scale-free to generate co-expression network modules among genes based on the topological overlap measure (TOM) with default parameters. Transcriptional regulatory networks were generated by assessing Pearson correlation coefficient (PCC > 0.85) between genes and TFs and by predicting *cis*- element binding sites in the promoter regions of key genes within the same module. The networks were visualized using Cytoscape (Kohl et al., 2010) .

### Homologous genes related to trichome development

Our team assembled a high-quality genome of an autotetraploid *A. argyi*. Based on homologous alignment with Arabidopsis and *A. annua*, we identified the genes and TFs involved in trichome development in *A. argyi*. Then we mapped the trichome development pathway based on the literature and showed the expression heatmap in different tissues. We also conducted the association analysis of TFs related to the development of trichome in *A. argyi*.

### Phylogenetic tree of MIXTA/MIXTA-like proteins among multispecies

To explore evolutionary relationships of *MIXTA*/*MIXTA-like* genes, 144 MIXTA/MIXTA-like homologous proteins were downloaded from NCBI and Phytozome according to literatures (Bedon et al., 2014; Brockington et al., 2013) , including 136 Angiosperm, two Gymnosperm, and six ANA grade as outgroup species. We constructed ML (maximum likelihood) tree using IQ-TREE and iTOL (https://itol.embl.de/) was used to visualize and edit the tree (Minh et al., 2020) . Meanwhile, there are three types of angiosperms including 21 monocots, 114 dicots, and one magnoliid species.

### Cloning and heterologous transformation of *AarMIXTA1.2*

Total RNA of *A. argyi* was reverse transcribed to cDNA using a PrimeScript RT Reagent Kit (TaKaRa, China). The primers were designed for PCR of candidate genes (Table S2). Each 10 μL reaction mixture of RT-qPCR contained 4.4 μL of cDNA template, 0.3 μL of each primer, and 5 μL of 2 × SYBR Green Pro Taq HS Premix (Accurate, China). The expression value for each gene was calculated by the 2−ΔΔCt method. The data presented were the mean ± SD of transcript expression levels obtained from at least three independent experiments. Then the target gene was sequentially connected to intermediate vector Blunt and PCAMBIA2300 vectors to obtain PCAMBIA2300-Blunt-AarMIXTA1.2 plasmid. The constructed plasmid was transformed into Agrobacterium, and Arabidopsis was infected using floral dip method and waited for maturity to harvest. After sowing, Arabidopsis was screened for positive plants. RT-qPCR was performed using SYBR Green Pro Taq HS premix qPCR kit according to the instructions. The 2–ΔΔCt method was used to calculate relative expression of genes. The method of phenotypic observation of transgenic Arabidopsis was similar as “Preparation and observation of *A. argyi* for microscope”.

## Results

### Trichome characteristic of *A. argyi* leaves with microscopes

Trichomes as the main tissues of moxa floss are the specific characteristics of *A. argyi*. Fluorescence microscope and scanning electron microscope were used to observe trichomes of the upper and lower epidermis to understand their morphology and distribution. The upper epidermis of *A. argyi* leaves was dark green, and the lower epidermis was slightly lighter in color, showing gray-green (Fig. S1). Through scanning electron microscope, it can be clearly seen that there are a large number of GSTs on the upper epidermis, and TSTs were relatively sparse (Fig. 1a). TSTs are densely entangled and distributed in the lower epidermis, and GSTs can be seen sporadically. The morphology of GSTs and TSTs can be seen under both scanning electron microscope and fluorescence microscope. The GSTs were in the shape of shoe soles, and the two arms of the TSTs are of unequal length.

**Figure 1.**
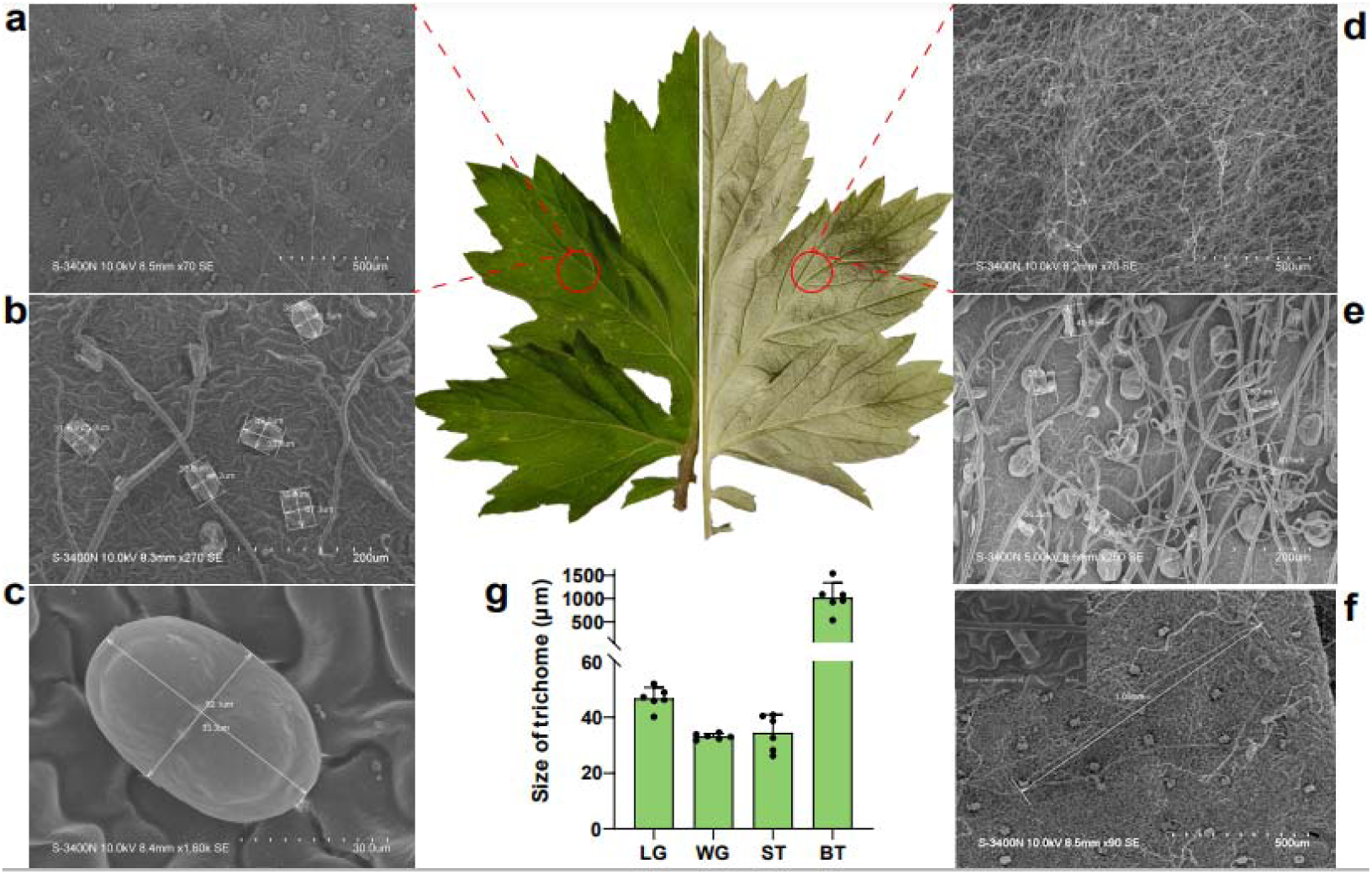
Characteristic of trichomes of *A. argyi* leaves using scanning electron microscope. a, b, c: GSTs of upper epidermis of A. argyi leaves at 8.5 mm × 70 SE, 8.3 mm × 270 SE and 8.4 mm × 1.6 k SE. d, e: TSTs of lower epidermis of *A. argyi* leaves at 8.2 mm × 70 SE, 8.6 mm × 250 SE. f: The branching length of the TST at 8.5 mm × 90 SE. The upper left corner of the image showed the stalk length of the TST at 8.6 mm × 1.3 k SE. g. The size of trichomes on upper surface in *A. argyi*. LG: length of GSTs; WG: width of GSTs; ST; stalk length of TSTs; BT: branching length of TSTs.

Two types of trichomes unevenly scattering on leaves of *A. argyi* were observed with scanning electron microscope. The GSTs with 33.3 μm × 52.1 μm and TSTs with 28.8 μm stalk length and more than 1 mm branching length were obviously visible on the upper epidermis in mature leaves (Fig. 1b, c, e, f), and their numbers and density were obviously higher than that of Arabidopsis and *A. annua* (Fig. S2). While what is special is that the overlong TSTs are distributed in high density on the lower epidermis of the leaves and intertwined together, making it difficult to distinguishing them during measurement (Fig. 1d, e, f).

### Expression differences between TSTs and non-TSTs of *A. argyi* leaves

For understanding regulation mechanism of abundant TSTs in *A. argyi*, TSTs were manual separated from the lower epidermis of leaves for RNA-seq (Fig. 2a). A total of 39.03 Gb CleanBase was obtained from transcriptome sequencing, with the effective data volume of each sample ranging from 6.27 to 6.88 Gb. Reads were aligned to *A. argyi* genome, with an alignment rate from 74.30% to 97.15% (Table S3, S4). The three biological replicates of the two tissues have a good clustering relationship (Fig. 2b). There were 8,901 up-regulated genes and 4,698 down-regulated genes were identified in TSTs compared non-TSTs tissues (Fig. 2c, d). The up-regulated genes of integral component of membrane, plasma membrane, defense response was significantly enriched, which were mainly related to development and function of TSTs (Fig. 2e, f). More R2R3-MYB proteins, C2H2 zinc-finger protein, homeodomain-leucine zipper (HD-Zip) IV protein, basic helix-loop-helix (bHLH) domain-containing protein, ethylene-responsive TF, and WD40 repeat-like superfamily protein (Yang & Ye, 2013; Chalvin et al., 2020) , which were considered as play important roles in regulation of trichome development, were up-regulated expression in TSTs. KEGG enrichment analysis showed that pathways of differentially expressed genes (DEGs) mainly included plant MAPK signaling pathway, isoflavonoid, flavonoid, etc. This showed that differences between TSTs and non-TSTs are concentrated in the synthesis and metabolism of substances, signal transduction, etc., which may be related to the fact that GSTs can synthesize, store and secrete secondary metabolites.

**Figure 2.**
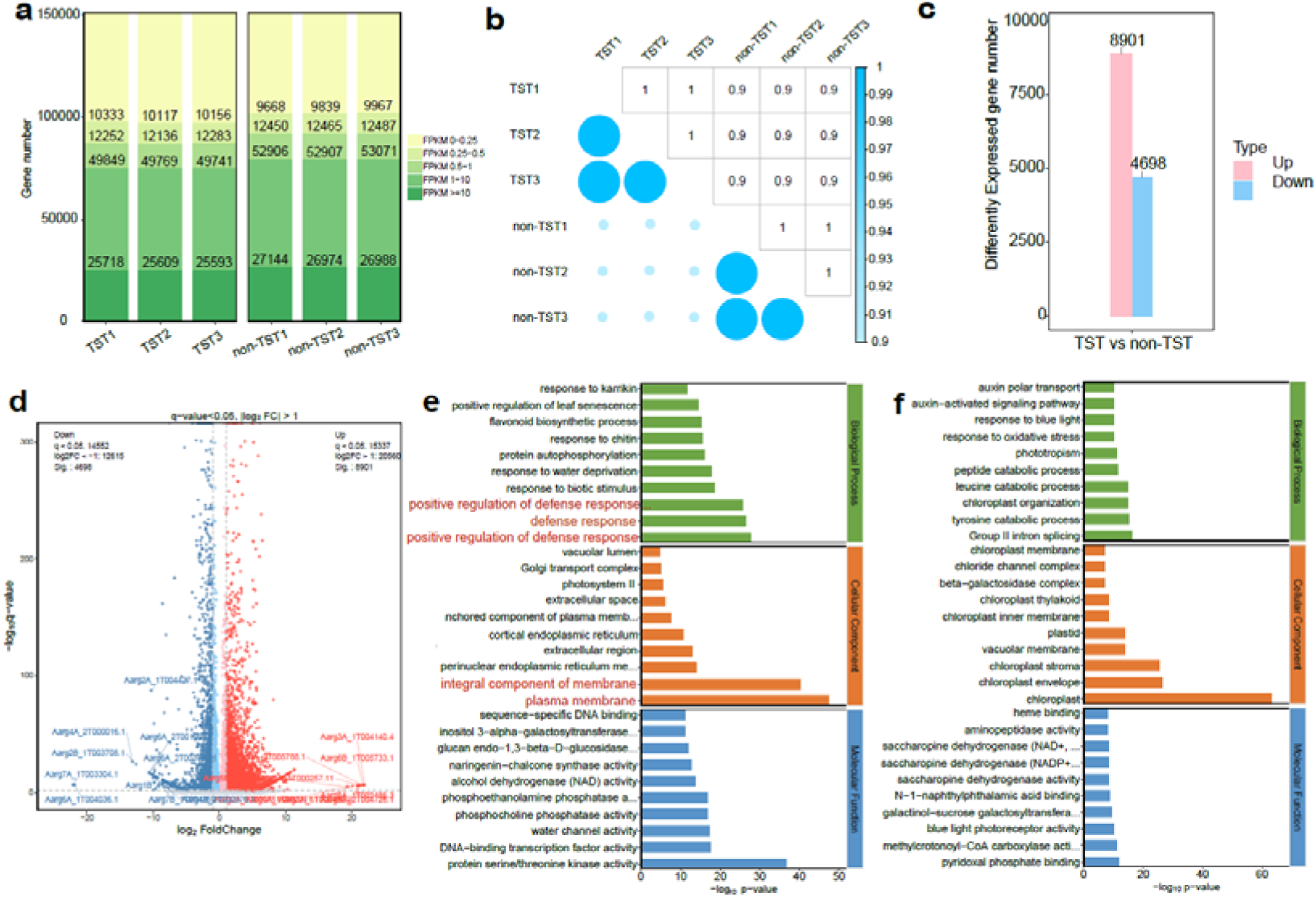
Gene expression of TST and non-TST in *A. argyi*. a: The FPKM ratio of total genes. b: The correlation of sequenced samples. c: The numbers of DEGs between TSTs and non-TSTs. Value parameter: *q*-value < 0.05 and |log2FC| > 1. d: Volcano plot of DEGs between TSTs and non-TSTs. e: Top 30 GO term of up-expression genes of TSTs vs non-TSTs. f: Top 30 GO term of down-expression genes of TSTs vs non-TSTs.

### Transcription factor and homologous genes analysis

To clarify TFs involved in trichome development regulation, we integrated transcriptome data of old leaves (OL), young leaves (YL), stem (St), and GST in NCBI, TSTs and non-TSTs, and a scale-free WGCNA network was constructed using the 53,354 genes (TPM ≥ 5), resulting in the identification of five distinct modules (Fig. 3a). Module cyan (MEcyan) was significantly enriched in the cutin biosynthetic process and contained homologous genes involved in Arabidopsis trichome development, including *GRA*, *TT8*, *MIXTA*, *GL2*, and *HD1* (Fig. 3b). Further visualization of the co-expression relationships among TFs within MEcyan revealed *AarMIXTA*s as Hub genes were significantly related with other regulatory TFs (MYBs, bHLH, WD40, HD-ZIP, WRKY, etc.) in correlation regulatory network (Fig. 3d).

**Figure 3.**
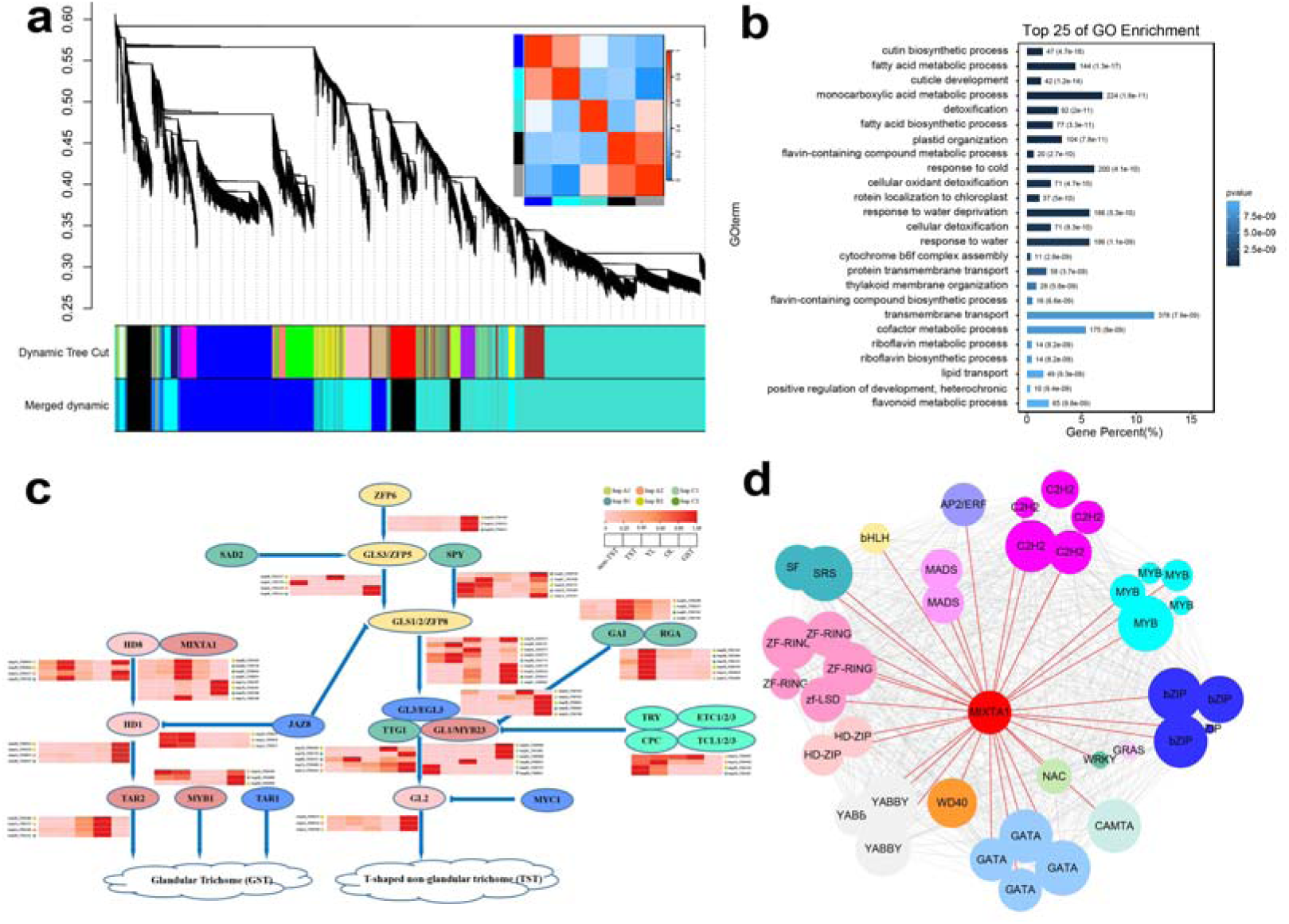
WGCNA of genes in *A. argyi*. a: Module partitioning, clustering and correlation between modules. b: GO term of Module cyan. c: Identification and expression analysis of homologous TFs regulating glandular development in *A. argyi*. d: WGCNA of trichome regulatory TFs in *A. argyi*.

Considering that certain key TFs involving in trichome formation has been reported in Arabidopsis and *A. annua* (Yan et al., 2018; Yang & Ye, 2013; Chalvin et al., 2020) . Based on homologous alignment, 119 genes belonged to 27 allelic groups involving in trichome development were identified in *A. argyi* (Table S5). Among them, 14 groups (*MYB*, *HD*, *GIS*, *ZFP* etc.,) maintain four allelic genes (ohnologs) and six groups (*GL1*, *GL2* etc.,) possess three allelic genes with one genes deletion. *ETC1*, *TT8* and *EGL3* showed tandem amplification, and *TTG1*, *RGA* and *MIXTA*s showed allelic amplification. *MIXTA*s possessed eight genes in *A. argyi* genome (Table S5). These trichome development genes have relatively significant tissue-specific expression (Fig. 3c). The major allelic genes among homologous chromosomes mentioned above showed similar expression trend, such as *ZFP6*, *HD1*, *TAR2*, *JAZ8*, *GAI*, and *RGA*. However, other genes (*MIXTA1*, *HD8*, *GLS1*/*2*/*ZFP8*, etc.) showed expression differentiation (Fig. 3c). Meanwhile, there are eight *MIXTA* genes in *A. argyi*, which is much more than that of Arabidopsis and *A. annua*, we speculate *MIXTA* genes have an important influence on the formation characteristics of mugwort trichomes.

### Phylogenetic analysis of multispecies *MIXTA*/*MIXTA-like* genes

To explore evolutionary relationships of *MIXTA*/*MIXTA-like* genes, we constructed an ML phylogenetic tree (Fig. 4a). 144 MIXTA/MIXTA-like protein sequences were downloaded from NCBI and Phytozome according to literatures (Brockington et al., 2013; Bedon et al., 2014) (Fig. 4a, Table S7). The tree was rich in species, including 136 Angiosperm, two Gymnosperm, six ANA grade (outgroups) species. There are three types of angiosperms: 21 monocots, 114 dicots, and one magnoliid species. Except for six outgroups, the remaining members were divided into four clades from I to IV (Fig. 4a). The MIXTA/MIXTA-like protein numbers from clades I to IV were two, 26, 33 and 77, respectively. Clade I had the two Gymnosperm species, indicating an earlier origin. Clade II included 21 monocots, four dicots, and one magnoliid species that differentiated earliest. Clades III and IV only included dicots, suggesting that the two clades may originated from clades I and II, and MIXTA/MIXTA-like proteins in clades I and II appeared before the divergence between monocots and dicots. Judging from the distribution of MIXTA/MIXTA-like of the same species in the tree, most species acquired ancestral MIXTA/MIXTA-like genes and then were duplicated or doubled within species in evolution. For example, MIXTA/MIXTA-like proteins of Asteraceae, Brassicaceae, Fabaceae, Malvaceae, Ranunculaceae, and Solanaceae showed species clustering. The evolution of MIXTA/MIXTA-like proteins in each family of the tree is basically consistent with APG IV system (https://www.mobot.org/MOBOT/research/APweb/).

**Figure 4.**
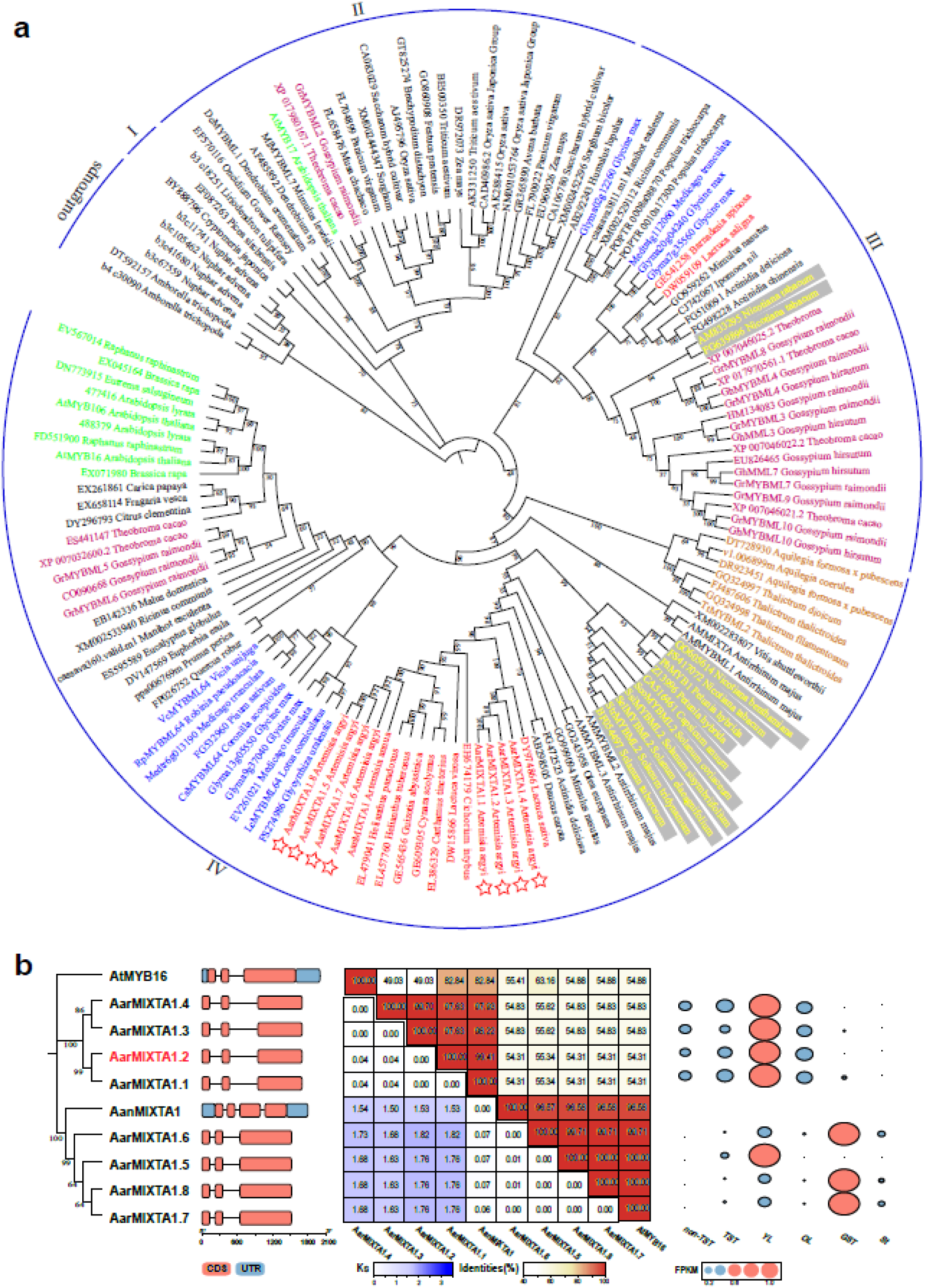
Phylogenetic analysis. a. The ML tree of MIXTA/MIXTA-like proteins of multispecies. Red, green, blue, purple, orange, and yellow fonts represent species from Asteraceae, Brassicaceae, Fabaceae, Malvaceae, Ranunculaceae, and Solanaceae families. Red stars indicate eight MIXTA proteins of A. argyi. b: Relationship, gene structure, similarity, and differential expression in various tissues of *AarMIXTA* allelic genes.

Eight *AarMIXTA* genes were found in *A. argyi* through homologous alignment, indicating the duplication compared with *A. annua*. They were divided into two clusters in the ML tree. AarMIXTA1.1, 1.2, 1.3, 1.4 were clustered with *Lactuca sativa*, and the other four AarMIXTAs were clustered with *A. annua*. The duplication of *AarMIXTA*s occurred after the differentiation of Asteraceae from other families (Fig. 4a). Higher *Ks* between the two clusters of *AarMIXTA* indicated that the two group of genes had a high mutation rate within *A. argyi*, which may due to the region where these genes are located experiencing a high recombination rate, less selection pressure, or positive selection in some populations (Fig. 4b). Higher *Ks* values may also be associated with gene duplication events, such as the expansion of *MIXTA*s. The allelic *AarMIXTA* genes had identical gene structure and sequence similarity with *AanMIXTA*s (Fig. 4b and Fig. S5). However, there was obviously expression difference in allelic *AarMIXTA*s (Fig. 4b). The expression of cluster of *AarMIXTA1.1*, *1.2*, *1.3*, and *1.4* in TSTs was higher than that of cluster of *AarMIXTA1.5*, *1.6*, *1.7*, and *1.8*, while the expression of cluster of *AarMIXTA1.5*, *1.6*, *1.7*, and *1.8* in GSTs was much higher than that of cluster of *AarMIXTA1.1*, *1.2*, *1.3*, and *1.4*, further illustrating the functional differentiation of *AarMIXTA*s (Fig. 4b).

### *In vivo* assay of *AarMIXTA1.2* transgenic Arabidopsis

*AanMIXTA1* in *A. annua* has been verified to positively regulate the density of GSTs, has been verified *in vivo*, so the function of its cluster (AarMIXTA1.5, 1.6, 1.7, 1.8) may be more clear. The function of the other cluster (AarMIXTA1.1, 1.2, 1.3, 1.4) is more likely different to that of *AanMIXTA1*. So *AarMIXTA1.2* were selected from the cluster for *in vivo* verification. The expression of *AarMIXTA1.2* were up-regulated in accordance with leaf developments and reached highest expression in third leaves. *AarMIXTA1.2* were specific up-expression in TSTs than non-TSTs based on qRT-PCR (Fig. 5a, Table S7). The *AarMIXTA1.2* were further cloned and transformed into Arabidopsis (Fig. 5b). It was found that compared with wild-type Arabidopsis (Col-0), the number of leaves of transgenic Arabidopsis (pCAMBIA2300-35S::*AarMIXTA1.2*) increased significantly, and trichomes were denser (Fig. 5c). In order to explore the changes in the density of trichome in transgenic Arabidopsis, their leaves were observed using stereomicroscope and scanning electron microscope. The density of TSTs, branch length, stalk length and other characteristics were statistically analyzed and calculated. Compared with wild-type Arabidopsis, TSTs density of transgenic Arabidopsis increased by about one time, and the branch length and stalk length increased slightly (Fig. 5d).

**Figure 5.**
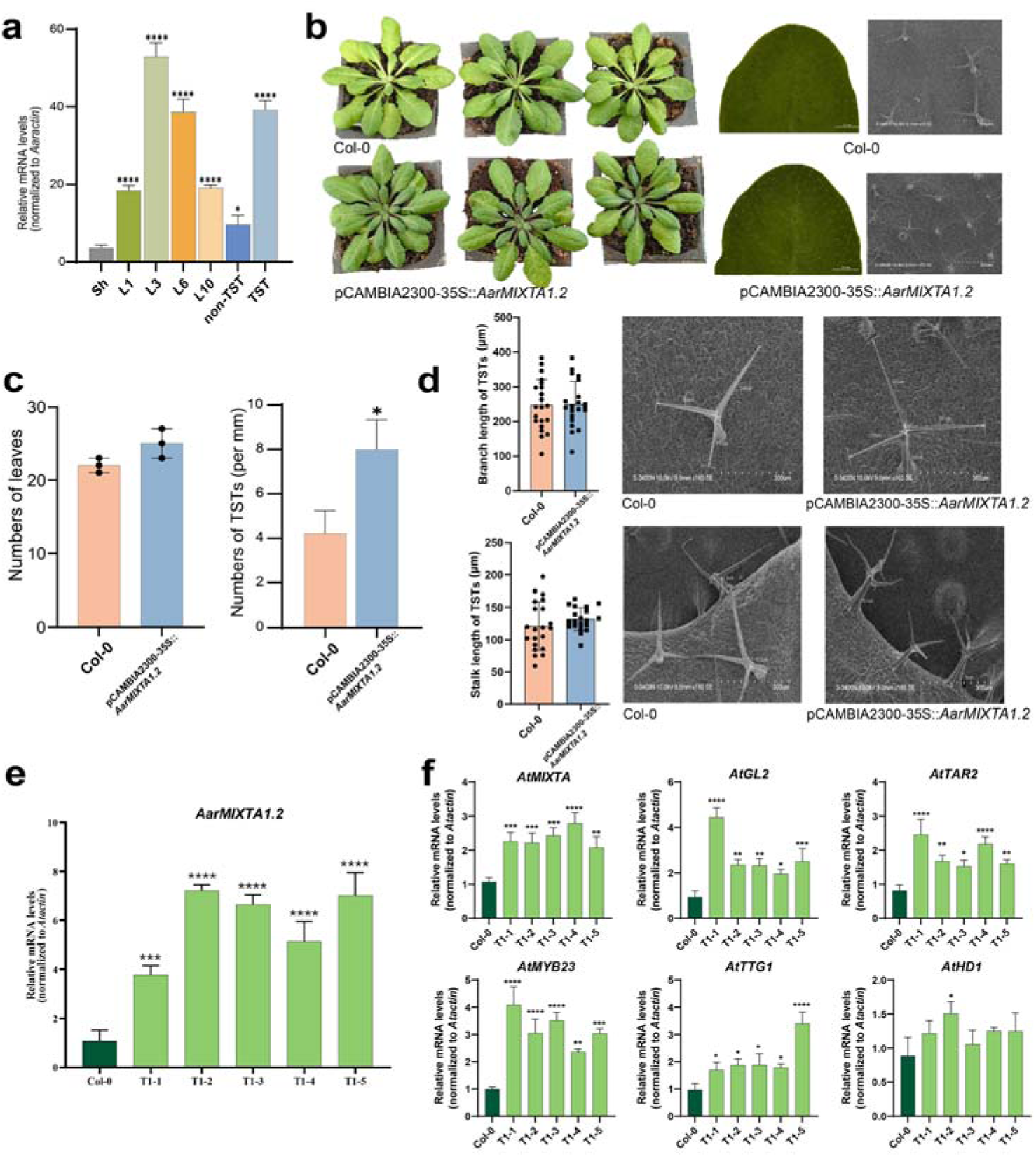
The transgenic Arabidopsis overexpression *AarMIXTA1.2*. a: Expression trend of *AarMIXTA1.2* in different leaf developments, TSTs and non-TSTs. Sh, L1, L3, L6, L10 represented leaf buds, the first group, the third group, the sixth group, the tenth group of leaves, respectively. b: The seeding, leaves and its upper epidermis with scanning electron microscope of Col-0 and over-expression pCAMBIA2300- 35S::*AarMIXTA1.2* Arabidopsis. c: Changes in leaves numbers and TSTs density in transgenic Arabidopsis leaves. d: The change of branch and stalk length in transgenic Arabidopsis leaves. e: Gene expression of over-expression pCAMBIA2300- 35S::*AarMIXTA1.2* Arabidopsis using qRT-PCR. f: The expression leaves of key TFs responding to *AarMIXTA1.2* in Arabidopsis using qRT-PCR.

Further, qRT-PCR was used to detect transcription level of *AarMIXTA1.2* in overexpressed *AarMIXTA1.2* Arabidopsis. The expression levels of *AarMIXTA1.2* in transgenic Arabidopsis were higher than those of wild-type Arabidopsis, indicating the *AarMIXTA1.2* transgenic plants were successful (Fig. 5e). To understand the molecular mechanism in overexpressed AarMIXTA1.2 plants, the expression levels of key genes *AtGL2*, *AtTAR2*, *AtMYB23*, *AtTTG1*, and *AtHD1* as downstream regulation factors in trichome development pathway were also detected. Their expression levels in transgenic Arabidopsis were significantly increased (Fig. 5f). At the same time, the original *AtMIXTA* in Arabidopsis also were promoted to express (Fig. 5f). The above results indicated that over-expression of *AarMIXTA1.2* may positively regulate TSTs development by regulating the downstream genes, which also showed that *AarMIXTA1.2* played an important regulatory role in the trichome development.

## Discussion

The significant difference in the distribution of GSTs and TSTs on the upper and lower epidermis of *A. argyi* leaves makes it an ideal system for studying the development of multicellular TSTs. Although the developmental regulation mechanism of single-cell TSTs has been studied in Arabidopsis and cotton, the mechanism involving in high density and different upper and lower epidermis of multicellular TSTs is more complex and unclearly. Our research started with microscopic observation of trichome of *A. argyi*, comparison of transcriptional expression of TSTs and non-TSTs, phylogenetic analysis of *MIXTA*/*MIXTA-like*, and experimental verification of *AarMIXTA*, and explained the intrinsic reasons for the development of TSTs in *A. argyi* in a hierarchical manner.

MIXTAs, belong to subgroup 9 (SBG9) R2R3 MYB TFs, which were important for specification and regulation of plant cellular differentiation (Brockington et al., 2013) . Many species with distinctive trichomes have *MIXTA*-related studies. *AanMIXTA1*, a regulator of cuticle biosynthesis, promote GST initiation to enhance artemisinin content in *A. annua* (Shi et al., 2018) . *CsMIXTA* involved in glandular trichome morphogenesis in *Cannabis sativa* (Haiden et al., 2022) . The high density of overlong TSTs on the lower epidermis of leaves is classical feature of *A. argyi*. The expansion and functional differentiation of *AarMIXTA*s are the characteristics of *A. argyi* MIXTA. The resulting gene expression dosage effect and neofunctionalization make TSTs of *A. argyi* much richer than that of Arabidopsis and *A. annua*. *AarMIXTA1.2* overexpression obviously increased the density of TSTs, which showed functional differentiation with *AanMIXTA*s to some extent. Meanwhile, most TFs showed allelic genes amplification and tandem repeat amplification, with some allelic genes also displaying deletions or becoming pseudogenes after genome duplication. Among them, *AarMIXTA*s as core genes showed balanced allelic amplification and result in the functional differentiation after genome doubling.

Of course, this study can potentially be expanded in a lot of areas. *A. argyi* has a strong adapt ability to the environment in China. Whether this ability is related to the development of its trichomes requires more sample collection and comparison. The mechanism between *AarMIXTA*s and the corresponding proteins and their chromosome accessibility is also worth further study.

In conclusion, we experimentally confirmed the role of *AarMIXTA1.2* in boosting TSTs formation and performed an evolutionary analysis of *MIXTA*/*MIXTA-like* genes. *AarMIXTA* genes have significantly expanded, which could be the underlying cause of its high TSTs in *A. argyi*. This research will greatly aid in enhancing the quality and yield of *A. argyi* as well as the development of novel cultivars through trait observation, gene screening, evolution analysis, and functional verification of TSTs.

## Supporting information

This table includes all the appendices marked in the text.

## Acknowledgments

This work was supported by National Natural Science Foundation of China Grant (Grant NO. U22A20446), Science and Technology Projects in Guangzhou (2023A04J0466) and Key Talent Attraction and Training Project (A1-2601-24-414- 112Z67-2024).

## Supplementary materials

Supplementary material associated with this article can be found, in the online version, at https:XXX.

---

**FigS. 1.**
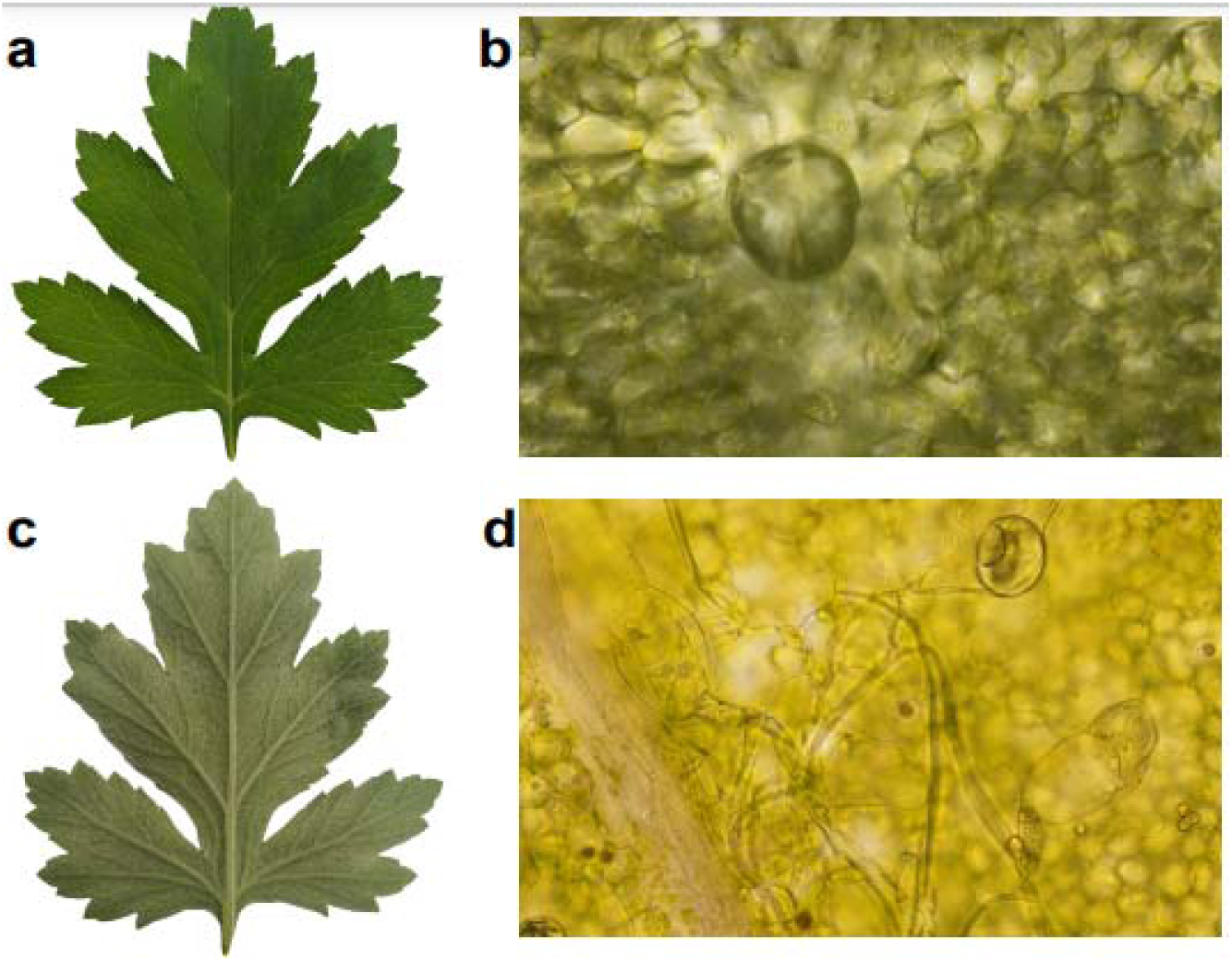
Microscopic observation of the epidermis of A. argyi. a: Upper epidermis. b: upper epidermis (Bright, fluorescence microscope ×40). c: Lower epidermis. d: Lower epidermis (Bright, fluorescence microscope ×40).

**FigS. 2.**
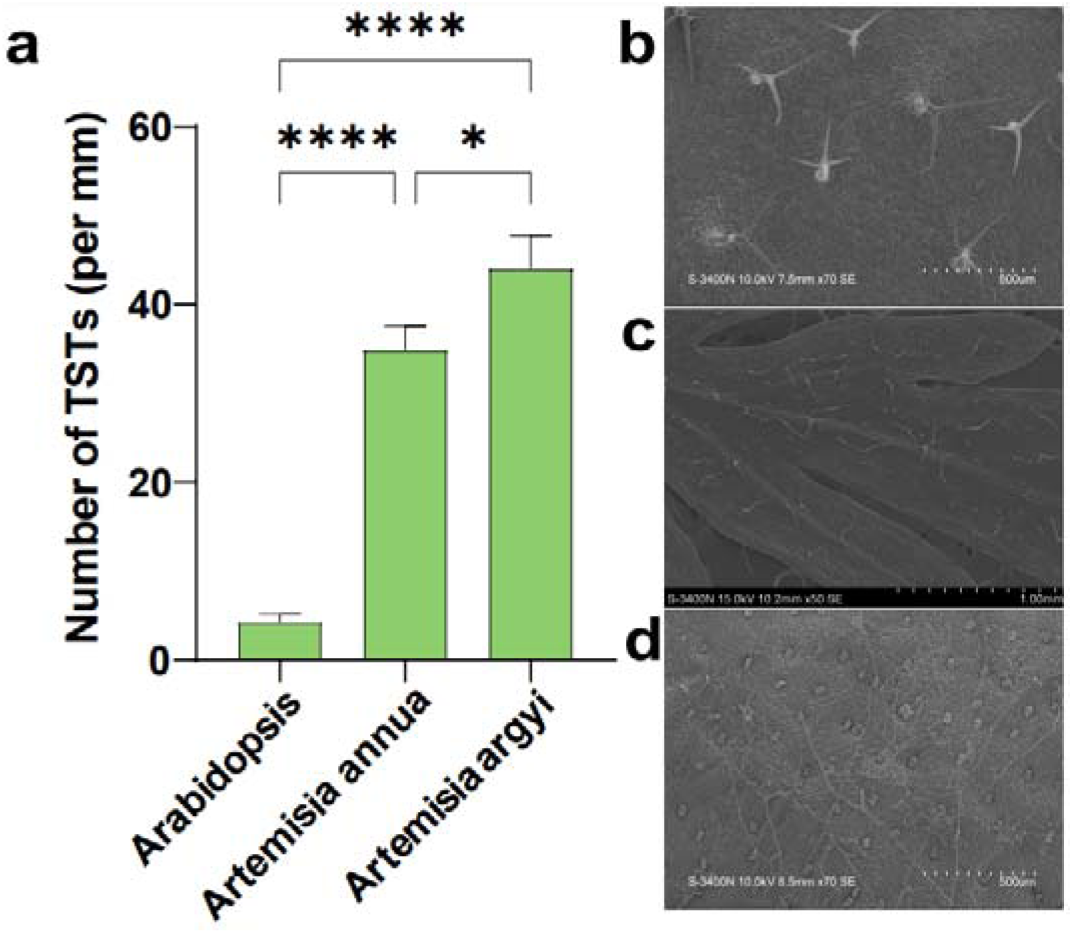
The density of TSTs on upper epidermis (a) and characteristics in (b) Arabidopsis, (c) *A. annua* and (d) *A. argyi*.

**FigS. 3.**
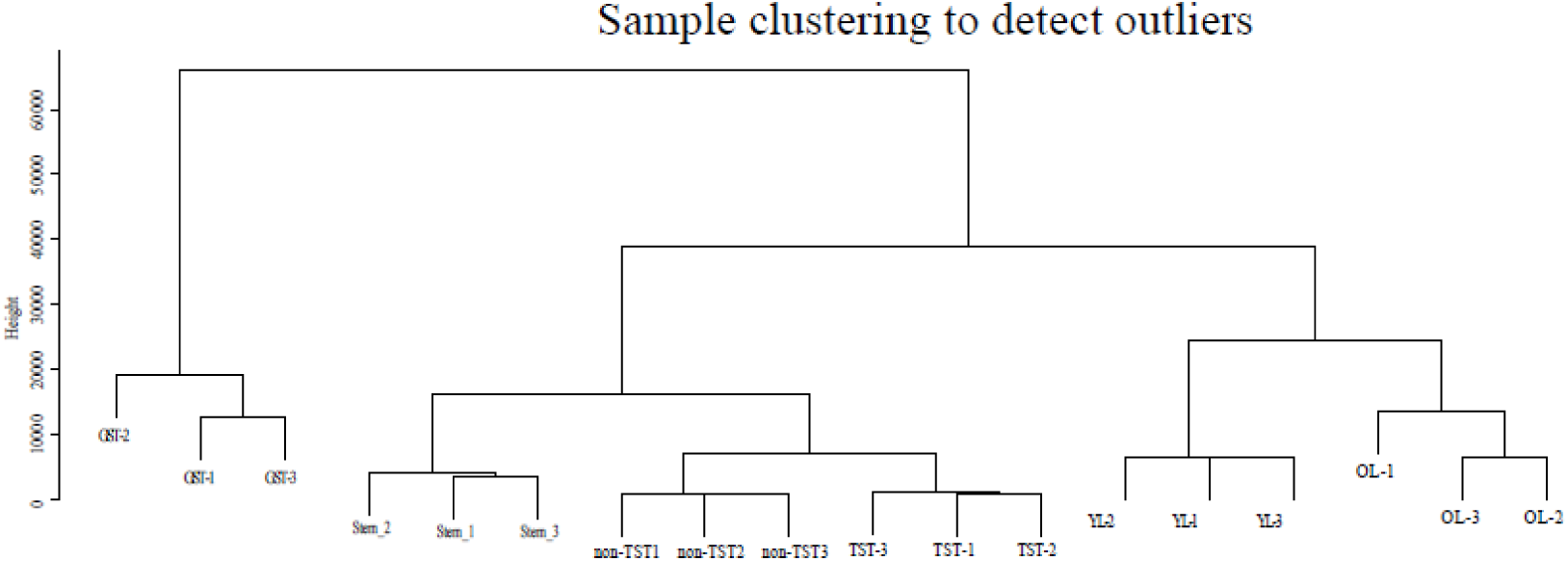
Sample clustering. GST: Trichome (NCBI); St: Stem; non-TST: mesophyll; TST: Trichome (ours); YL: Young_leave; OL: Old_leave.

**FigS. 4.**
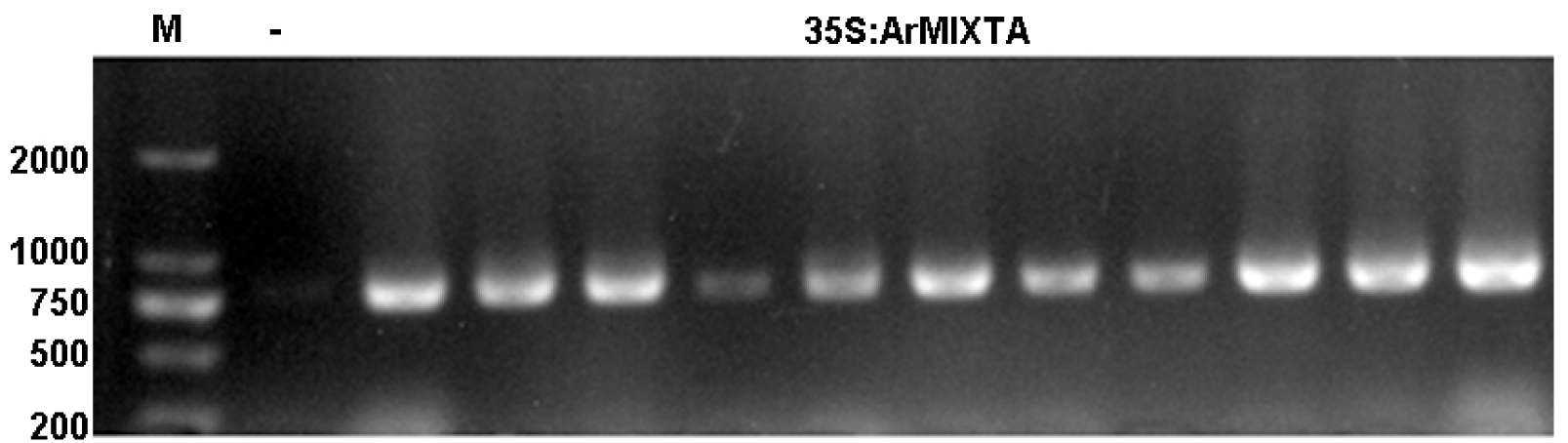
PCR verification of transgenic 35S::AarMIXTA Arabidopsis.

**FigS. 5.**
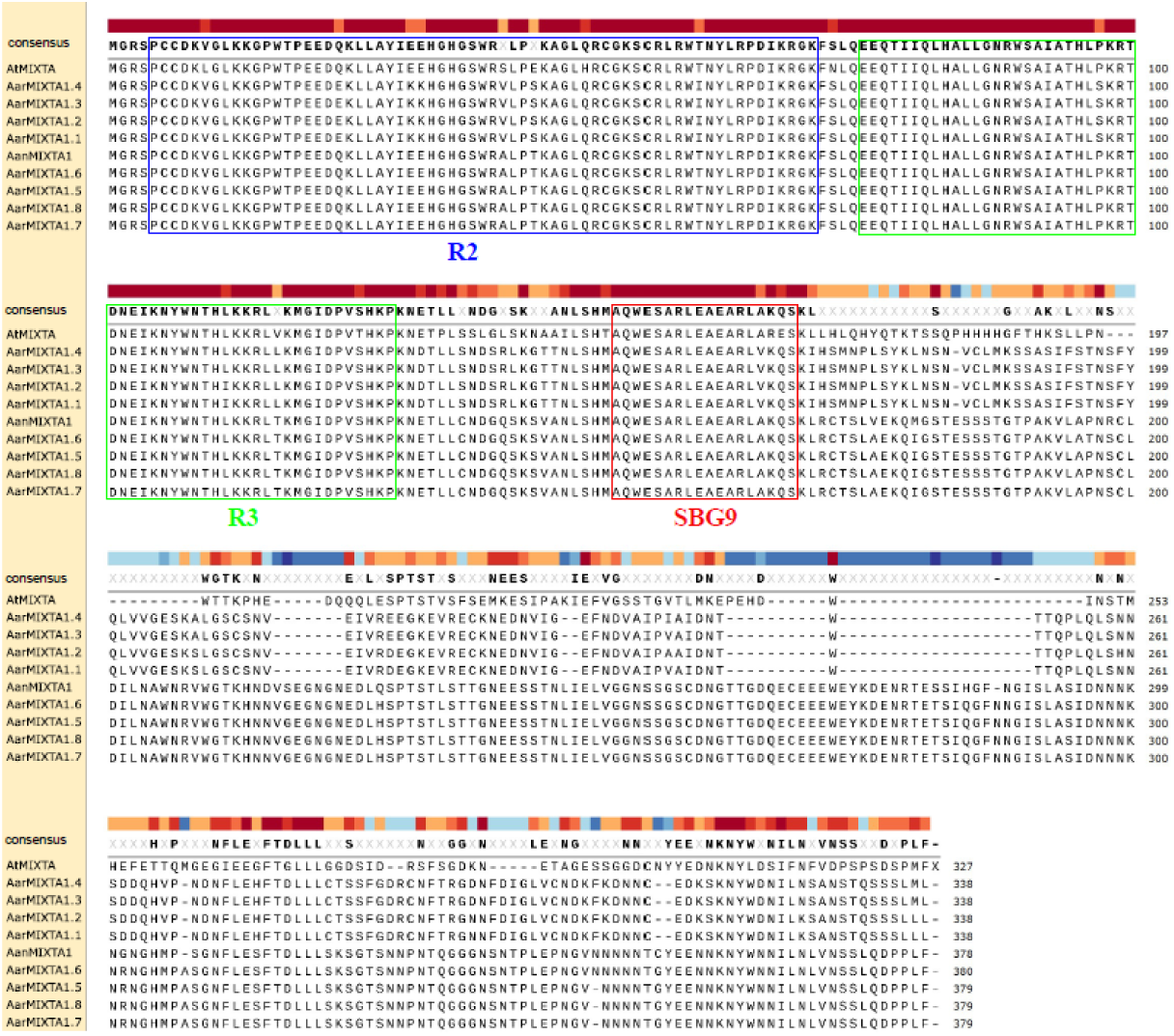
Sequence similarity comparison of MIXTAs in Arabidopsis, *A. annua* and *A. argyi*.

